# Spatial heterogeneity enhance robustness of large multi-species ecosystems

**DOI:** 10.1101/2021.03.23.436582

**Authors:** Susanne Pettersson, Martin Nilsson Jacobi

## Abstract

Understanding ecosystem stability and functioning is a long-standing goal in theoretical ecology, with one of the main tools being dynamical modelling of species abundances. With the help of dynamical population models limits to stability and regions of various ecosystem dynamics have been extensively mapped in terms of diversity (number of species), types of interactions, interaction strengths, varying interaction networks (for example plant-pollinator, food-web) and varying structures of these networks. Although it is apparent that ecosystems reside in and are affected by a spatial environment, local differences (spatial heterogeneity) is often excluded from studies mapping stability boundaries under the assumption of an average and equal amount of interaction for all individuals of a species. Here we show that extending the classic dynamical Generalised-Lotka-Volterra model into a connected space the boundaries of stability change. When viewing the ecosystem as a spatially heterogeneous whole, limits previously marking the end of stability can now be crossed without any remarkable change in species abundances and without loss of stability. Thus limits previously thought to mark catastrophic transitions are not critical due to the possibility of spatial heterogeneity within the system. In addition, we show that too much spatial fragmentation of ecosystem habitats acts destabilising and leads back to the stability boundaries found in spatially homogeneous ecosystems with average interactions. Thus, we conclude that spatially heterogeneous but connected systems are the most robust. In terms of ecosystem management, the risk of collapse or irreversible changes is lower in spatially heterogeneous systems, which real ecosystems are, and we should expect local changes in populations well in advance of system collapse. Although, too much fragmentation of an ecosystem’s available space can lead to a less robust system with higher risk of extinctions and collapse.

**Author summary:** One of the major challenges facing humanity is the fragmentation of wildlife habitats and decline in biodiversity due to human need for resources and land-use practices. We need to find ways to combine human prosperity with biodiversity conservation. To achieve this a solid understanding of ecosystem stability and functioning is paramount. One way to gain such insight is to find limits when we expect species to go extinct or ecosystems to collapse by simulations of interacting species populations. Many such stability limits have been found theoretically the last decades, but for simplification of modelling, studies often exclude that ecosystems are spread out in space. By doing so, studies assume an average and equal amount of interaction for all individuals of a species. Here, we explicitly include space and thus allow for migration and spatial heterogeneity (local differences) in interactions. When modelling the ecosystem as a spatially heterogeneous whole, limits previously marking extinction or collapse can now be crossed without any remarkable change in species abundances. Thus, natural hindrances to migration improve ecosystem robustness and limits previously thought to mark catastrophic transitions are not critical due to the possibility of spatial heterogeneity within the system. In addition, we reconfirm that a large amount of fragmentation acts destabilising for an ecosystem. Thus, we conclude that the most robust ecosystems are spatially heterogeneous but connected.

## Introduction

There are many ways to think about and represent ecosystem functioning and stability, one prominent direction in mathematical ecology is the search for limits of stability in dynamical species population models in terms of some specified parameters or characteristics of the system. To this end Generalised-Lotka-Volterra (GLV) dynamics and modifications, with for example higher order interaction [1] (ex. third species modifying how one species interacts with another) as well as many examples of more advanced interaction functions between species have been used. With a proliferation of stability results both confusing and enlightening, such as the role of diversity for stabilising ecosystems [2–4] or to what extent properties such as modularity [5] or nestedness [6] in interactions act stabilising or destabilising or are aspects of the same property [7].

Specifically the question whether diversity (number of species) and/or strength of interaction between species act destabilising or not, has drawn a lot of attention since the first mathematical formulations of large ecosystem stability with May’s paper in the 1970:ties [2]. The conclusions are not uni-vocal but a majority of studies find that at a certain level of diversity (or interaction strengths) the system will no longer have a stable equilibrium solution [8]. Meaning that ecosystems cannot sustain too high diversity or interaction strength and can possibly collapse if interaction strengths are increased when for example an ecosystem’s available space is reduced. Studies have also located feasibility (all species non-extinct) limits where higher diversity or interaction strength lead to extinctions (loss of feasibility) [9, 10], and systems vulnerable to perturbations in structural features (such as growth rates, or interaction strengths) [11, 12]. With even higher diversity or interaction strength, abundances can start oscillating or the system collapses to a significantly smaller subset of species [9]. This latter limit is the classical limit recognised as a drastic change in system characteristics and many times referred to as collapse.

A prominent model indicating diversity as destabilising is the original Generalised-Lotka-Volterra model. One of the many simplifications of the GLV and other extended models is the lack of space. This simplification leans on the assumption that if two species interact, all individuals in the two species interact the same amount. An assumption that no real ecosystem fulfils. Nevertheless, many of the conclusions on stability are drawn from models without the spacial dimension.

There are on the other hand studies specifically taking space into account, studying for example pattern formation where the diversity is small [13–15], space influence on interaction structure [16], spread of specific species [17] or meta-community studies of different flavours, of local stability in the feasible domain [18], synchronisation of single species populations [19, 20] or exhibiting macro-ecological patterns at the edge of collapse [21]. A feature that still holds in the latter type of spatial stability analysis is that there is a boundary of radical shift and loss of stability. However, there are no studies mapping all stability boundaries and behaviours in large spatial multi-species ecosystem.

In this paper we drop the assumption of average interaction for the entire ecosystem by simulating large multi-species ecosystems spread out over a connected space. This allows us to re-investigate stability boundaries, ecosystem responses to pressures and the effect of ecosystem fragmentation.

## Theory and methods

The version of the classical GLV model we will use as a base is stated below

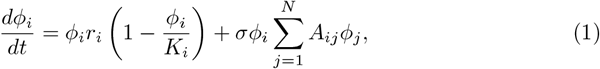

where *ϕ*_*i*_, *r*_*i*_ and *K*_*i*_ are species abundances, intrinsic growth rates and carrying capacities for species *i* respectively. The web of interaction between species is represented by *A*_*ij*_, a *N × N* matrix with a density of interaction of *c* = 0.5 and whose entries are drawn from a normal distribution with negative mean, *µ* = *−*0.5 and a variance of one. This means all types of interactions (trophic, mutualistic, competitive etc.) are included although there is a larger percentage of competitive (-,-) and amensialistic (-,0) interactions. The parameter *σ* becomes the standard deviation (s.d.) of the interaction strengths and is often used as a tuning parameter and proxy for complexity, an increase in interaction strengths means an increase in complexity.

For small *σ* in systems governed by Eq 1 with diversity *N* the system will settle in a unique stable fixed point with all *N* species present. When increasing *σ* (increasing complexity) at a certain value of *σ* feasibility is lost, meaning to stay in a stable fixed point species will successively go extinct if increasing *σ*. Further increase in eventually pushes the system across the final stability boundary and the system transitions to either oscillations, chaos or a fixed point with a substantial loss of species. This latter limit has many times been referred to as collapse. The region between the two boundaries is structurally unstable meaning a small perturbation in parameters (*r*_*i*_, *σ, c* etc.) lead to qualitative change, in effect species extinctions [9]. This region can also have multiple stable fixed points with differing patterns of non-extinct species [22].

We introduce a spatial dimension into the GLV model by setting up a grid (or line) where all grid-points have the same interaction matrix. This in effect, allows for the same maximum amount of species who interact in the same way at every grid-point. The grid-points are connected by diffusion, representing the migration of species in space. This can be seen as for example animals migrating to areas where there is a larger abundance of prey per predator or plants dispersing to areas where there are less competitors (more available resources).

To avoid effects from the boundary of the grid we use periodic boundary conditions (shape the grid as a torus in two-dimensional space or ring in one-dimensional space) as shown in Fig 1.

**Fig 1.**
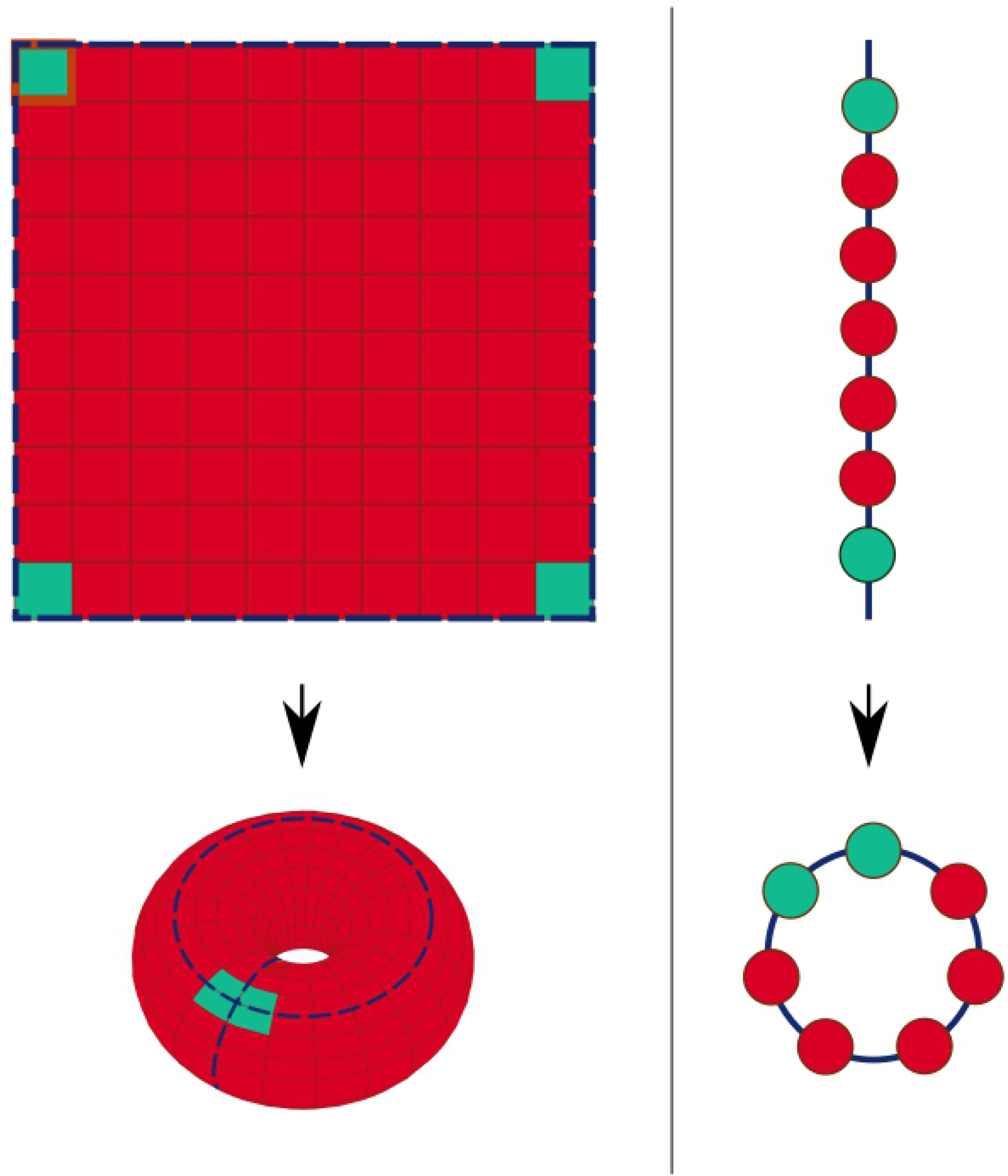
Schematic grid and line. The figure shows schematic pictures of how the grid is formed like a torus (to the right) and how the line is formed like a ring (to the left). This is done to simulate an ecosystem situated in a large connected space and to minimise boundary effects.

The GLV equations in continuous space with diffusion are

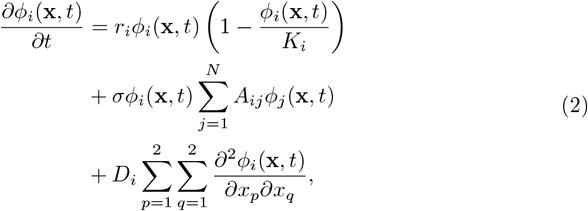

where *ϕ*(**x**, *t*)_*i*_ is the relative abundance of species *i* which now depends on both spatial location **x** and time *t*. The diffusion rates for species *i* are *D*_*i*_, which are different for all species but same for all gridpoints for the same species and are random variables drawn from a uniform distribution. Since we wish to use a grid we discretise the spatial dimension of Eq. 2 using the a discrete Laplace operator in two or one dimension given as

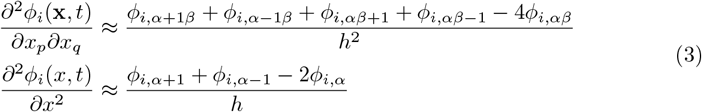

where *α* and *β* are grid indices and the denominator *h* is the “distance” between patches set to 1. The resulting dynamical equation for species *i* in grid-point (*α, β*) in a two dimensional grid is thus

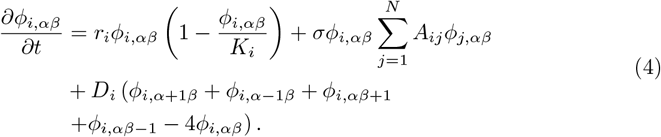

Real ecosystems are often situated in either two-or three-dimensional space. In most of our simulations on the other hand we used one-dimensional space to reduce the computational load and effect of boundaries. This was acceptable since we noticed no qualitative difference in results between two- and one-dimensional simulations.

## Results

With the addition of a connected space we add the possibility of different dynamics or solutions and timing of dynamics in different grid-points. Local heterogeneity (local species abundance differences) because of different dynamics is only possible when *σ* is large enough (larger than the feasibility boundary) so that different solutions of the equations are available in the grid-points and the system can end up with multiple fixed points, oscillatory patterns or a combination of fixed points and oscillations. The same oscillatory dynamics can also give rise to spatial heterogeneity by phase-shifts between grid-points. The amount of spatial heterogeneity depends on the magnitude of the diffusion constants *D*_*i*_. If diffusion constants are zero this represents a number of identical disconnected systems with GLV dynamics, while very large constants lead to complete synchronisation of the grid, smoothing out all local differences in the abundances of species. All these local variations lead to a multitude of system characteristics in terms of combinations of grid-point dynamics and phase-shifts.

### Local abundance oscillations and crossing stability limits

Species abundance oscillations can appear in the entire non-feasible region. We find that in many systems for the spatial GLV, increasing *σ* can make a fixed points transition to a oscillatory pattern with the same number of species. This behaviour is not as prevalent in the non-spatial GLV, which instead commonly switches to another fixed point with a smaller number of species, in effect species go extinct if the system is perturbed or pressured. The connected space acts as a buffer and the system can stay stable with either the whole system’s species abundances oscillating if the grid-points are synchronised or “stay” in the fixed point by local unsynchronised oscillations. The spatial systems are therefore more robust.

If diffusion constants are small so that synchronisation does not take place, and systems are at interaction strengths nearing previous collapse boundaries this means averaging over the local variations results in species abundances in accordance with fixed points that are unstable in the non-spatial GLV. Thus, this robustness of the spatial GLV makes it possible for the ecosystems to stay stable in parameter regions beyond stability limits of the non-spatial GLV.

In Fig 2 is an example of a system of *N* = 20 maximum biodiversity for a range of *σ* larger than the feasibility limit, and diffusion magnitudes leading to out of phase oscillations. We see that the unsynchronised local species abundance oscillations lead to stable species abundances of the whole ecosystem at an interaction strength where there is no fixed point in the non-spatial GLV. Thus the previous stability boundary is passed by the spatial GLV almost without any change of the system species abundances.

**Fig 2.**
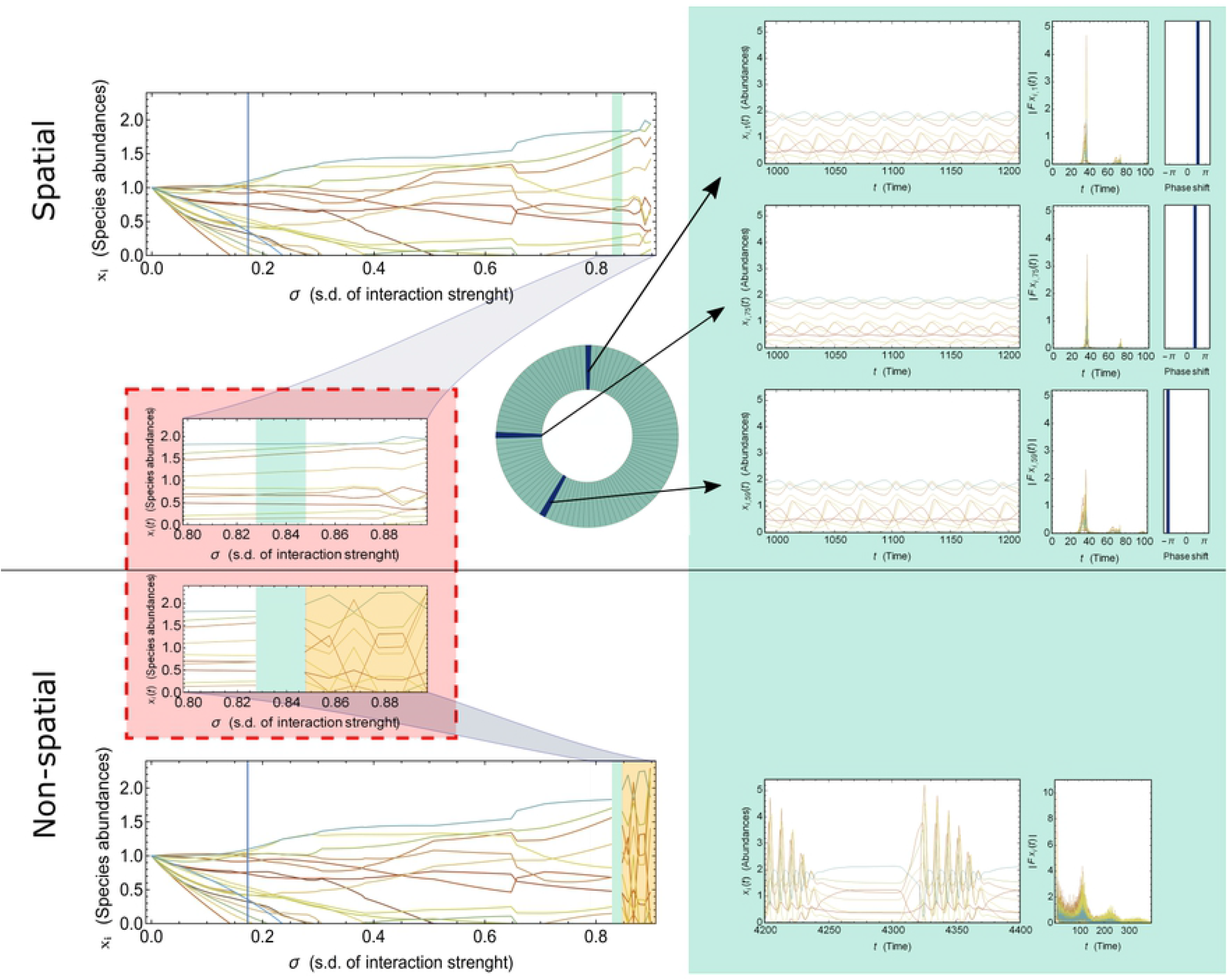
Unsynchronised local oscillations stabilise. The figure shows an example of a system of *N* = 20 maximum number of species with out-of-phase oscillations when *σ* is larger than the feasibility limit and diffusion rates are small. On the left side on top we show fixed point or oscillation grid-point average species abundances for a system with small diffusion and below fixed point abundances for the same system with no spatial dimension (non-spatial GLV). Both the abundances for the spatial system with diffusion and the non-spatial are shown for *σ* ranging from zero to collapse values with an enlargement of the latter part in the red box. The Green shading in the plots indicate a region where the only stable fixed point for the non-spatial version is with 3 species going extinct. On the other hand the abundance means in the spatial version show no change at all (green area in top left plots). With higher *σ* in the orange region of the non-spatial system (bottom left plot) the system is seen to be structurally unstable, while the spatial system on top shows little if any structural instability. This system behaves almost the same with and without a connected space until *σ* is large enough approaching collapse values. To the right on the green background are shown example dynamics for *σ* in the green marked area in the left plots, in different grid-points for the spatial system (top) and for a oscillatory solution for the non-spatial system (bottom). We see in the spatial system oscillatory dynamics in each grid-point example with the same Fourier spectra, but differing phases (the panels to the right). Together the different phases and amplitudes but same frequencies of the abundance oscillations average to the values corresponding to an unstable fixed point of the non-spatial GLV system. For the non-spatial system there is a oscillatory pattern, note however the increase in sharpness in both frequency and amplitude as well as some species going extinct and reappearing, which is not a biologically realistic or stable solution for a ecological system.

### Lower Variability

There are different ways the inclusion of a connected space stabilises the species abundance oscillations making them less variable. The oscillations in the spatial systems usually reflect a pattern found in the non-spatial GLV, but with the addition of diffusion they become less violent with lower amplitudes in oscillations. Other systems with oscillations might have different “preferred” oscillations for the non-spatial and spatial GLV respectively. In these cases the non-spatial oscillation patterns are higher in amplitudes and sometimes with higher frequencies than in the spatial GLV, see for example the non-spatial GLV oscillations in Fig 2 for one version of such an oscillation pattern. Yet another possibility is that diffusion makes possible a different lower amplitude oscillation pattern not present in the non-spatial GLV, an example of this is shown in Fig 3.

**Fig 3.**
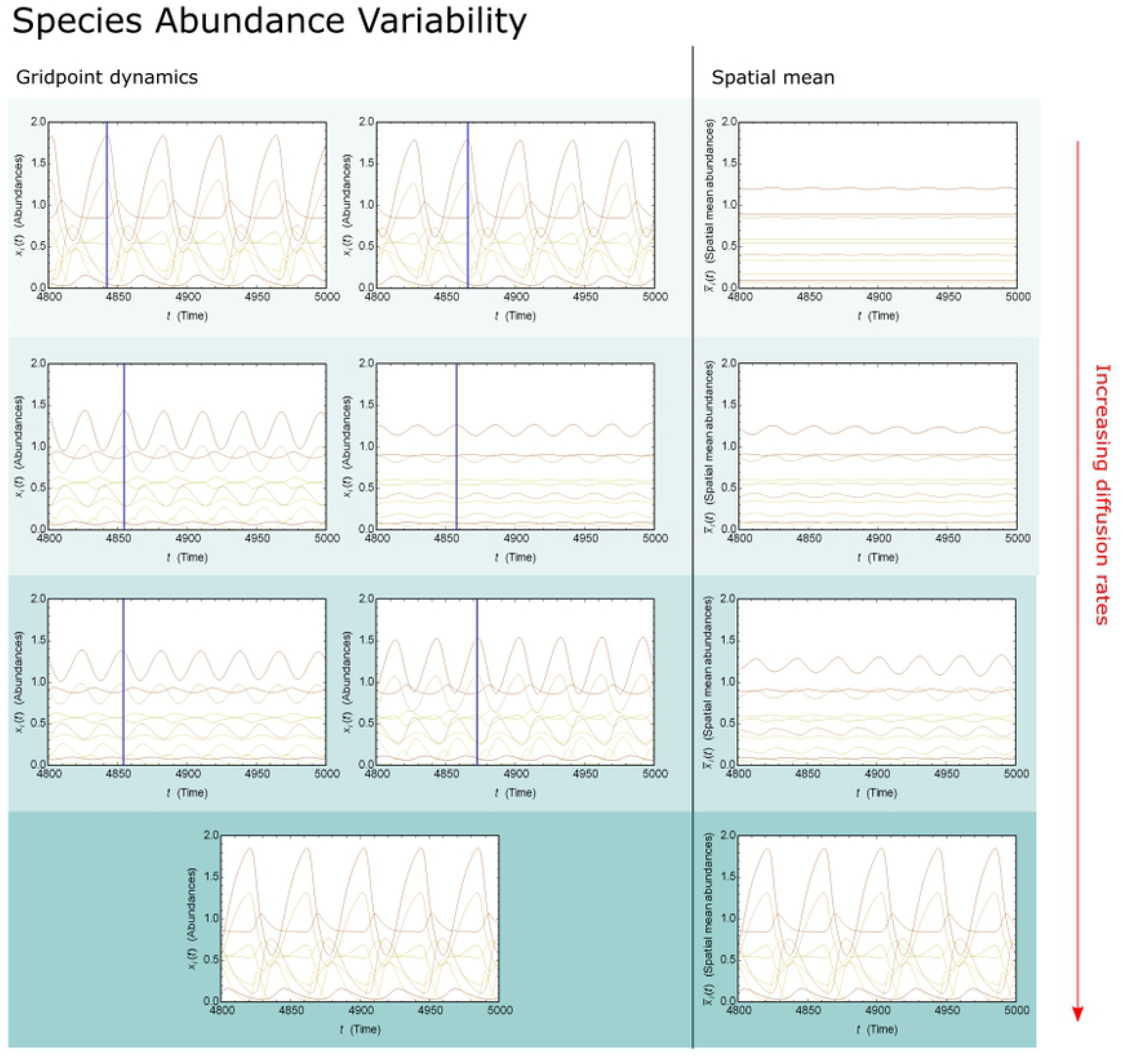
Species abundances in spatial extended system. The figure shows an example of a system with varying rates of diffusion (the green shading increases with increasing diffusion rates). The left column shows the system’s dynamics at arbitrary grid-points from high amplitude for the non-spatial system, as the amplitudes are dampened with increasing diffusion and then back to high amplitudes as the diffusion rate is high enough for synchronisation over the grid. If there are two panels in the left column this shows two unsynchronised grid-point dynamics. If there is only one panel the system is synchronised. The right column shows the spatial mean for all grid-points. Note that the unsynchronised oscillations lead to almost constant species abundances for the whole ecosystem while the synchronised system on the bottom shows violent oscillations for the whole ecosystem.

Many of these violent non-spatial oscillations lead to periodic species extinctions, an unrealistic scenario, while the spatial GLV avoids this both by less violent oscillations and by the connection to other local areas with the oscillation pattern shifted in phase. Significant lowering of the variability in species abundances we find when systems are unsynchronised at low to medium diffusion rates. Although such systems commonly have a large spread in oscillations amplitudes, all grid-points have dampened oscillation amplitudes compared to the non-spatial system and the abundances for the whole system are almost constant. When the diffusion rates in systems like these are increased, engendering synchronised oscillations, they usually reflect oscillation patterns present in the non-spatial GLV but again with reduced amplitudes.

In the limit of large *D* the ecosystems again retrieve their high amplitudes of the non-spatial GLV oscillations synchronised in all grid-points. Thus when diffusion is increased first oscillations start to synchronise leading to abundance oscillations for the whole ecosystem and with continued increase in diffusion the species oscillations increase in amplitude until they mimic that of the non-spatial GLV. It is therefore in the middle range of diffusion rates we find the most stabilising effects as seen in Fig 3.

### Phase-diagram

We have shown examples of dynamics of the spatial GLV and also pointed out the many possible ways ecosystems can behave in a connected space. We can bring some order to this multitude of behaviours and substantiate our claim that moderate diffusion rates lead to increased stability by gathering statistics on measures that contribute to stability, while varying *σ* and the diffusion rate. The measures we chose are 1) Average number of grid-points with oscillatory patterns (potentially different patterns), 2) Average phase shift between grid-points if oscillations (degree of synchronisation), 3) Average amplitude if oscillations, and 4) Average Diversity (*n*). The measures are both in combination and one-by-one the basis for different aspects of stability. Statistics over these measures in the non-feasible region for varying magnitude of diffusion rates are shown as phase diagrams in Fig 4.

**Fig 4.**
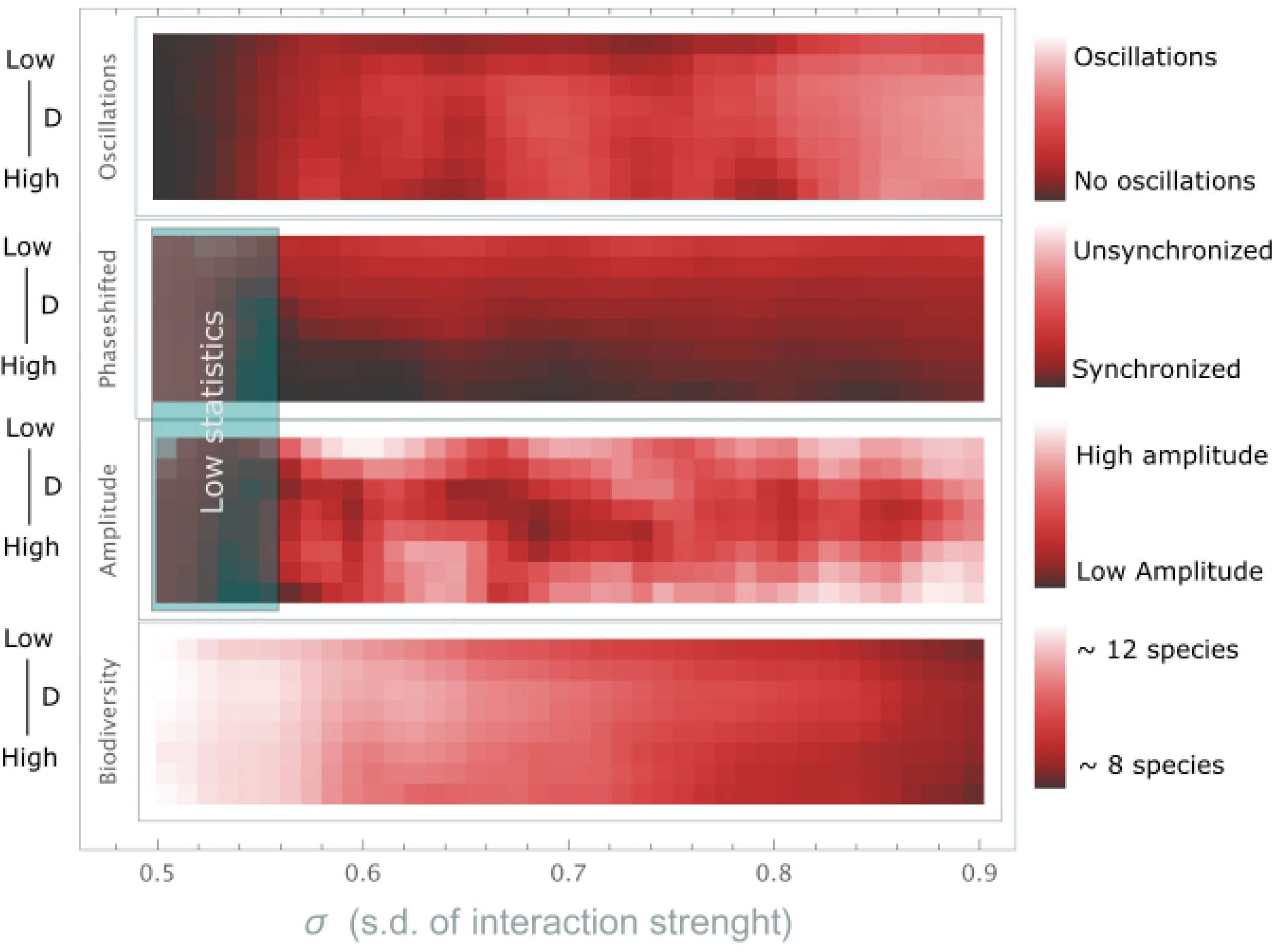
Phase diagram of diffusion magnitude vs. s.d. of interaction strength *σ*. The figure shows statistics in four phase diagrams of diffusion magnitude vs. standard deviation for 1) Average number of grid-points with oscillatory patterns, 2) Average phase shift between grid-points if oscillations (degree of synchronisation), 3) Average amplitude if oscillations, and 4) Average Diversity (*n*), from top to bottom respectively. The lowest diffusion in the diagrams is zero, corresponding to completely disconnected space (non-spatial GLV). Worthy of noting is the lower degree of oscillations and the lower diversity in the non-spatial systems. It is also clear that oscillations are present in almost the entire non-feasible region.

An example of the measures contributing to stability is, the oscillations appearing in almost the entire non-feasible region (although most prominently for higher interaction strengths) while the average diversity for systems with diffusion is higher than without diffusion. This shows the increase in robustness across the entire span of interaction strengths, *σ*. Systems can react with local oscillations instead of extinctions if pressured or perturbed. Another example, the lesser fraction of stable fixed points for large *σ* together with almost no oscillations for *D*_*i*_ = 0 (top row in diagram), show where systems without a connected space start collapsing. Adding moderate rates of diffusion so that the system has a connected space gives rise to out of phase oscillations and the ability for systems to stay stable across previous stability limits. The third diagram shows the decrease in species abundance variability (oscillation amplitudes) for the middle range of diffusion magnitudes, where both disconnected and completely connected lead to violent oscillations. This feature is connected to the degree of synchronisation shown in diagram two, where we can see the increase in synchronisation as the diffusion constants are increased in magnitude.

In Fig 5 we summarise the results from the phase-diagrams in terms of behaviour and stability. The table shows that the middle range of diffusion, in effect systems which allow for a certain amount of migration of species between local areas, are the more robust systems over a larger range of interaction strengths

**Fig 5.**
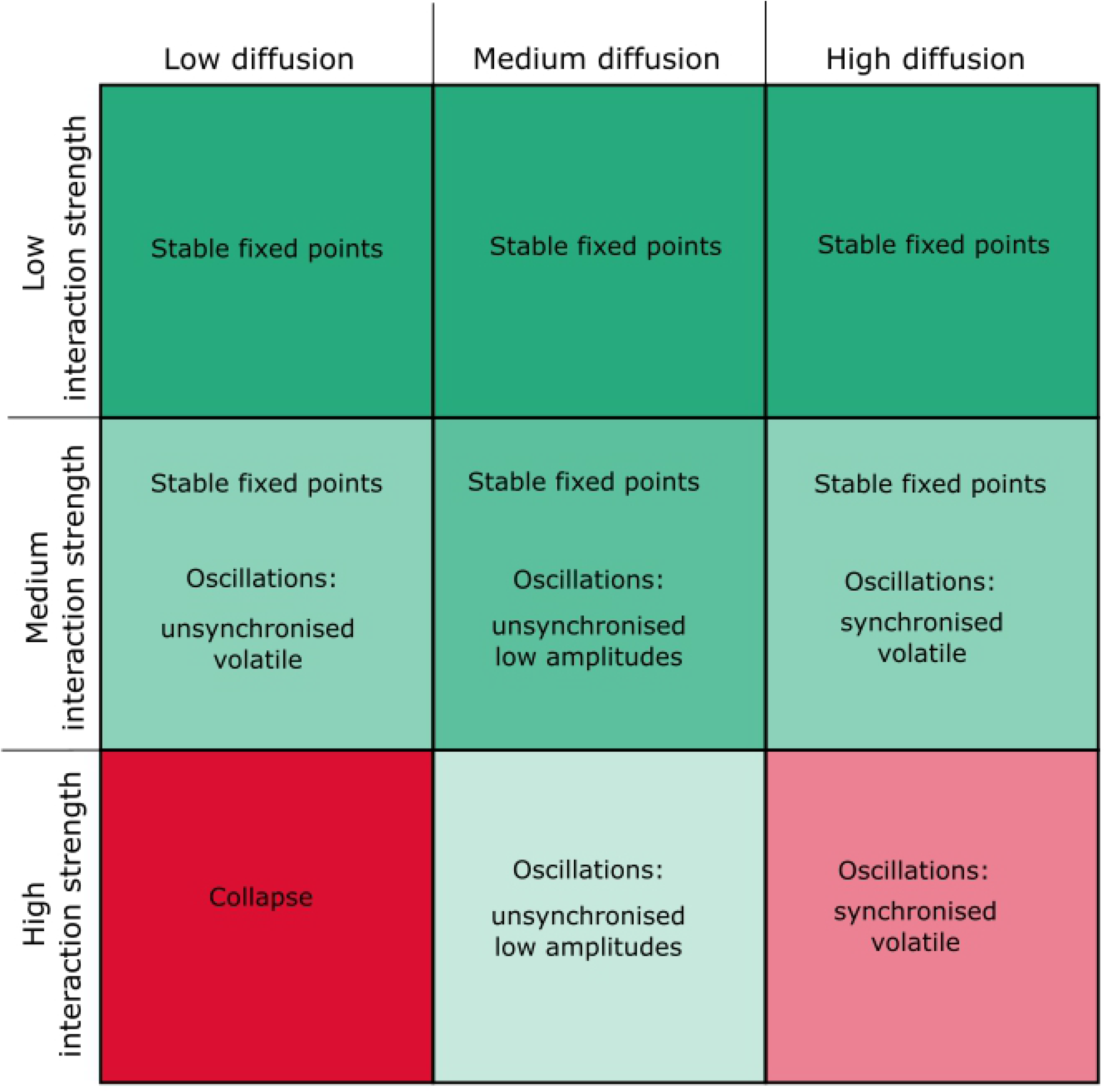
Stability of spatial GLV. The table shows in text the type of dynamics present for low, medium and high diffusion rates and s.d of interaction strength. Green indicates stable robust systems and the darker the colour the more robust. Red on the other hand are collapsed or unstable volatile systems. Note that the middle range is green for all s.d of interaction strengths.

## Discussion

Ever since the first mathematical model of an ecosystem in May’s paper [2], it has been an open question if the biodiversity, multitude of interactions between species and the amount of interactions (in effect complexity) of an ecosystem makes it more or less stable. Since then a limit to the complexity an ecosystem can sustain, derived in May’s paper and later extended, has persisted. Ecosystems at the edge of this limit or externally pushed so that their complexity exceeds it are predicted to radically change or collapse. Crucial aspects of such models are that the ecosystems are modelled as isolated homogeneous systems. A resolution to the long standing question of the limit of complexity might not be primarily the structure of interactions but that real ecosystems are both internally heterogeneous and externally connected. Connected ecosystems might gain their capacity of sustaining previously thought to be unstable patterns of cohabitation using the flexibility of local variations and buffer of its surrounding ecosystems.

As we have shown there are many aspects of stability that are affected when extending the GLV model to a connected space. Some effects are larger than others and put together they all act stabilising and blur out the boundaries of stability. Importantly we also find that ecosystems with a medium level of connection between local areas give the most stable and robust systems, a tendency also noticed in [18] and [**?**].

The classical stability boundaries have long been known to be a transition from a stable equilibrium fixed point to some qualitatively different dynamics in terms of abundance oscillations, chaos or drop to substantially smaller community [2, 9, 23]. Thus oscillations have been known to be a possible dynamics resulting from crossing the stability boundary. In a connected space the amplitudes of such oscillations are dampened. In addition, if the migration between local areas (diffusion rates) are low enough, the oscillations in species abundances can be out of phase leading to species populations as a whole remain largely unchanged. As real ecosystems are necessarily situated in space where animals interact with animals in their vicinity but occasionally migrate to nearby areas, intermediate diffusion rates in our connected space best approximates a real ecosystem. This would suggest that real ecosystem are less likely to abruptly transition to a qualitatively different state but will in advance experience stabilising local oscillations while keeping overall stability.

The effect of local stabilising oscillations is as we have shown above not only present when nearing collapse, but can also prevent extinctions when the system is perturbed. The non-spatial GLV is structurally unstable in the non-feasible region, meaning small structural changes (small increase in *σ*) can lead to species extinctions. The connected space, on the other hand can stabilise by local oscillations, thus avoiding the irreversible event of an extinction. With real ecosystems more alike the spatial GLV, we can expect ecosystem to be more robust than many models might suggest [24, 25].

We have argued that real ecosystems are in the middle range of diffusion, meaning in the middle range of migration. Thus, there is both a lower and higher limit for the system’s ability to buffer against perturbations. This suggests that ecosystem biodiversity and stability is not only sensitive to the known negative effects of fragmentation but also an increase in migration. This could for example result from the removal of natural boundaries. This is an effect not as known in ecosystem literature [26, 27] but the fact that similar GLV studies including a spatial dimension have found such models to reproduce known macro-ecological patterns (species abundance distributions, species area relations and range size distributions) [21] might lend some credence for the result. This might therefore be a real but less prevalent or hidden effect to take into consideration in ecosystem management.

The spatial GLV with ecosystems in a connected space can be further explored in many ways relevant to real ecosystems. A larger spread in diffusion constants for instance, can be used to better simulate different species mobility, since larger animals for example tend to move over larger areas. How this would effect stability is unclear although, since spatial patterning in reaction diffusion systems tend to appear in systems where the components have a large spread, we can speculate that species patterning might appear in such systems. The diffusion constants do not necessarily have to be equal between all grid-points either, varying constants can represent varying distance, some natural obstacle, a road or systems with internal dynamics connected to surrounding ecosystems. With varying diffusion constants we can therefore both study “unsymmetrical” habitats where interactions between local areas differ as well as how local disturbances might affect an ecosystem and connected ecosystems.

In line with studying the effects of local disturbances of diffusion rates, left for future work is varying the standard deviation of interaction strength *σ* over the grid. The inverse interaction strength can be interpreted as a proxy for available habitat, since a reduction of available space will force the existing animals closer together and to interact more which is an increase in *σ*. Thus, with different *σ* we can model a grid with different local habitat sizes as well as a system’s response to local habitat losses if perturbing one or some of the *σ*s at a time.

## Conclusion

Although modellers are well aware that real ecosystems are situated in space, the possible affects of spatial dimensions has rarely been accounted for in the GLV models. Many insights into ecosystems’ functioning have been gained with this simplification. However, this study explicitly shows that adding space to non-spatial dynamical populations models can significantly alter or modulate some of the conclusions. A limit of complexity for stability is no longer a limit to qualitative change or collapse, since the connected space acts as a buffer making the ecosystems more robust. Such systems in addition have lower variability and the ability to avoid extinctions by local species oscillations. We also find that systems with intermediate migration in the connected space are typically the more robust systems. Thus, both fragmentation of habitat or removal of natural boundaries can make an ecosystem more vulnerable to both extinctions and collapse.

